# Deep learning for supervised classification of temporal data in ecology

**DOI:** 10.1101/2020.09.14.296251

**Authors:** César Capinha, Ana Ceia-Hasse, Andrew M. Kramer, Christiaan Meijer

## Abstract

Temporal data is ubiquitous in ecology and ecologists often face the challenge of accurately differentiating these data into predefined classes, such as biological entities or ecological states. The usual approach consists of transforming the time series into user-defined features and then using these features as predictors in conventional statistical or machine learning models. Here we suggest the use of deep learning models as an alternative to this approach. Recent deep learning techniques can perform the classification directly from the time series, eliminating subjective and resource-consuming data transformation steps, and potentially improving classification results. We describe some of the deep learning architectures relevant for time series classification and show how these architectures and their hyper-parameters can be tested and used for the classification problems at hand. We illustrate the approach using three case studies from distinct ecological subdisciplines: *i*) insect species identification from wingbeat spectrograms; *ii*) species distribution modelling from climate time series and *iii*) the classification of phenological phases from continuous meteorological data. The deep learning approach delivered ecologically sensible and accurate classifications demonstrating its potential for wide applicability across subfields of ecology.

## 1. Introduction

The recent increase in affordability, capacity, and autonomy of sensor-based technologies (Bush et al., 2017; Peters et al., 2014), as well as an increasing number of contributions from citizen scientists and the establishment of international research networks (Bush et al., 2017; Hurlbert and Liang, 2012), is allowing an unprecedented access to time series of interest for ecological research. A common aim of ecologists using these data concerns assigning them into predefined classes, such as ecological states or biological entities. Typical examples include the recognition of bird species from sound recordings (e.g. Priyadarshani et al., 2020), the distinction between phases in the annual life cycle of plants (i.e., ‘phenophases’) from spectral time series (Melaas et al., 2013), or the recognition of behavioral states from animal movement data (Shamoun-Baranes et al., 2016). Many other examples exist, with scopes of application that range from the molecular level (Jaakkola et al., 2000) to the global scale (Schneider et al., 2010).

The assignment of time series into one of two or more predefined classes (hereafter referred to as ‘time series classification’; Keogh and Kasetty, 2003) can be performed using a variety of different approaches, ranging from manual, expert-based, classification (Priyadarshani et al., 2020) to fully automated procedures (see Bagnall et al., 2017 for examples). In ecology, time series classification is generally approached by processing the time series data into a set of summary variables − using hand-designed transformations, or techniques such as Fourier or wavelet transforms − and then using these variables as predictors in ‘classical’ supervised classification algorithms, such as logistic or multinomial regressions or random forests (e.g. Capinha, 2019; Dyderski et al., 2018; Priyadarshani et al., 2020; Reside et al., 2010; Shamoun-Baranes et al., 2016). In machine learning terminology, this approach is known as ‘feature-based’, where the ‘features’ are the variables that are extracted to summarize the time series.

Despite the wide adoption of feature-based approaches, important limitations still undermine their predictive performance and scalability. A key constraint concerns the need for domain-specific knowledge about the phenomenon that is being classified, to obtain ‘optimal’ sets of features with respect to predicting the reference classes. While this may not seem limiting, considering the ever-growing body of knowledge in the ecological literature, few, if any, ecological phenomena are fully understood (Currie, 2019). This inherently limits and casts doubt about the optimality of human-mediated selections of ‘relevant’ predictors. This limitation can be illustrated for species distribution modelling, a popular field among ecological modelers. These models often rely on readily available sets of predictors that summarize long-term climate averages and variability, (e.g. the ‘BIOCLIM’ variables; Booth et al., 2014), despite recognition that species distributions can also respond to short-term meteorological variation (e.g. Reside et al., 2010). Accordingly, these temporally aggregated predictors cannot guarantee a comprehensive representation of the role of climate in determining the distribution of species. Additionally, scaling modelling frameworks can result in reliance on pre-processed predictors because performing species-specific feature extraction could be prohibitively costly, in terms of human and time resources, when modelling the distribution of hundreds of species.

Here we focus on the use of deep learning models as an alternative to feature-based approaches for supervised classification of time series data. Deep learning models are a set of recent, complex architectures of artificial neural networks (Christin et al., 2019; LeCun et al., 2015), which have enabled significant advances of performance in highly complex tasks, such as computer vision and natural language processing (LeCun et al., 2015). In ecology, the number of studies applying these models is growing rapidly, but the vast majority of applications are to the classification of image and sound data (e.g. Brodrick et al., 2019; Campos-Taberner et al., 2020; Christin et al., 2019; Mac Aodha et al., 2018; Willi et al., 2019). Accordingly, we wish to draw the attention of ecologists to the capacity of deep learning models to classify all kinds of temporal data and to several of their practical and conceptual advantages in this regard.

A key difference between deep learning approaches and feature-based approaches for time series classification is that the former can perform the classification from raw time series data. For ecologists, this ability can be seen not merely as a methodological particularity, but as a conceptual and operational improvement from traditional modelling approaches. On one hand, the direct use of time series data as classification predictors positively forces ecologists to consider the temporal component of the analyzed phenomena (Ryo et al., 2019; Wolkovich et al., 2014), and, on the other hand, it relieves them from subjective decisions about the transformation of the temporal data. The identification of relevant features in the time series is performed automatically by the models and this procedure is guided by their capacity to accurately classifying the classes (Fawaz et al., 2019). Accordingly, a promise of these models is that they may capture relevant information (e.g. thresholds, lag effects; carryover effects; Ryo et al., 2019) that would be missed if relying on subjective sets of user-defined features, ultimately improving predictive performances. Additionally, because there is no need of human intervention in feature extraction, deep learning models allow a full, end-to-end, automation of computational workflows.

Below we provide a brief explanation of how supervised deep neural networks work and describe some of the modelling architectures more relevant in the context of time series classification. Next, we demonstrate the application of deep learning models as a general approach for time series classification using three case studies from distinct sub-fields of ecology. First, we perform species identification based on recordings of insect wing flap movements, second, we predict the potential distribution of a vulnerable mammal species using time series of monthly climate data, and third we predict the seasonal patterns of fruiting of a mushroom species, based on meteorological time series. We implement our models using a standardized modelling approach which should be accessible to the generality of ecological modelers, including non-experts in deep learning.

## 2. Material and Methods

### 2.1. Deep neural networks for time series classification

Artificial neural networks (ANN) are mathematical models inspired by how biological nervous systems process information. These models are often conceptualized in terms of nodes (or ‘neurons’) and weighted links. A basic ANN architecture includes a first layer of nodes, representing the input data, a second (‘hidden’) layer with nodes performing data aggregation followed by nonlinear transformation, and a final (‘output’) layer where the predicted values are computed (Fig. 1a). The nodes in each layer are connected to the nodes in the next layer through weighted links. The training of ANNs proceeds by iteratively adjusting the weights of links between the layers. An important notion is the ‘epoch’, which refers to when the entire training dataset is passed forward and backward across the network one time. During each epoch, the weights are updated to improve the network’s predictions, given the information fed to the input layer. For more details on ANNs with emphasis on ecology see, Lek and Guégan (1999), Olden et al. (2008) and Christin et al. (2019).

**Figure 1.**
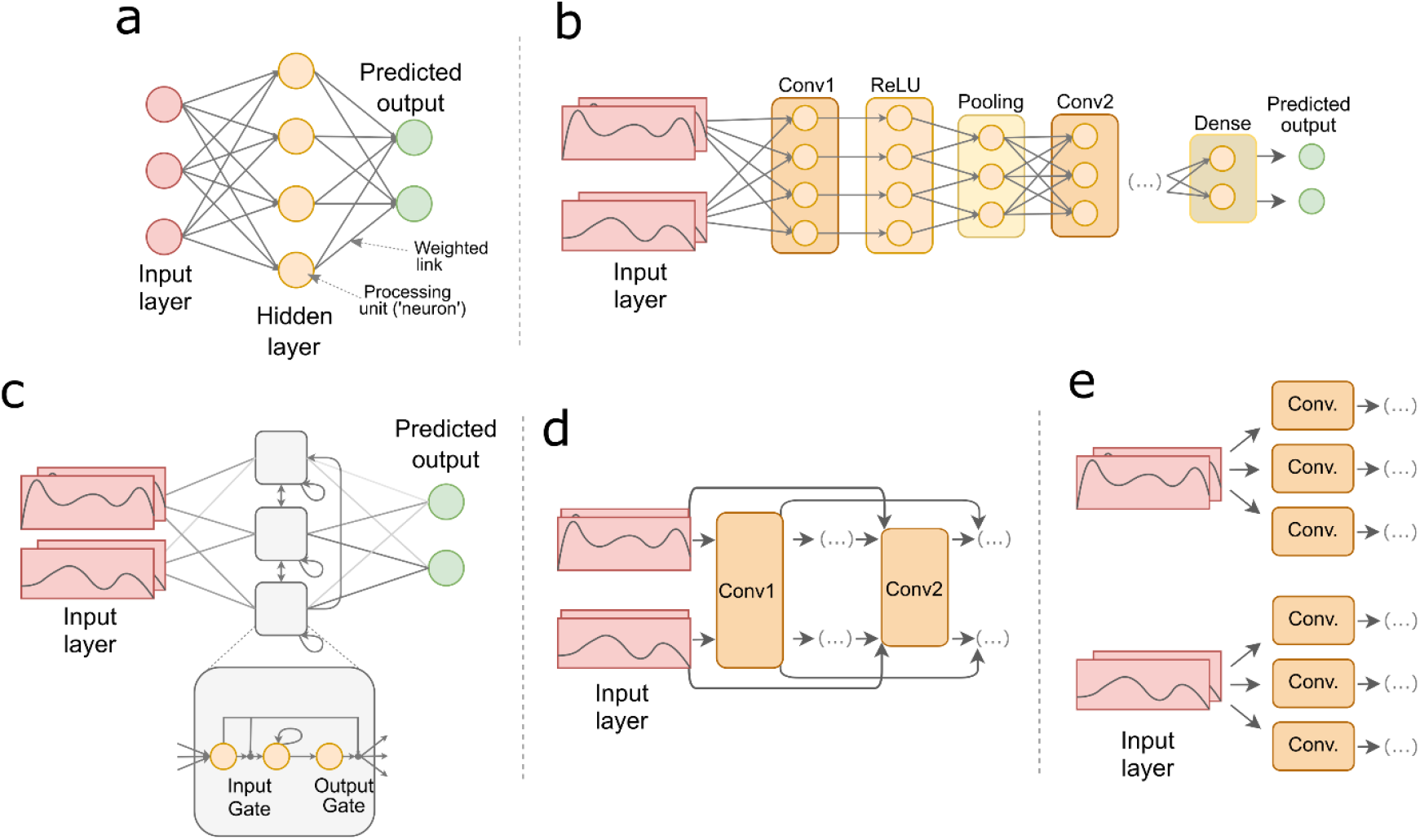
Schematic diagrams of artificial neural networks architectures and components. (a) A simple artificial neural network; (b) a classification Convolutional Neural Network; (c) a Recurrent Neural Network with Long Short-Term Memory units (grey squares); (d) residual blocks in a Convolutional Neural Network; (e) parallel stacking of convolution layers, as used in Inception Time Networks.

‘Deep’ neural networks refer broadly to ANN architectures that are capable of training large numbers of hidden layers and neurons (LeCun et al., 2015; Schmidhuber, 2015). This capacity determines the level of abstraction that the models can attain in representing the input data. Models with more hidden layers can capture more complex patterns and achieve a deeper hierarchy of features. In other words, shallow models tend to capture ‘basic’ patterns (e.g. a ‘spike’ in a specific time step), while deeper models are able to ‘learn’ more complex abstractions (e.g. spikes combined with a reduced long-term variability).

Many deep learning architectures can be used for time series classification (Fawaz et al., 2019; Wang et al., 2017). These architectures differ in the number of layers, and the mathematical functions the layers perform, as well as in the way information flows between them. Below we provide a description of four architectures used for time series classification. These architectures were chosen because they are widely adopted for time series classification and because they are available in mcfly (the software we use here for model implementation; van Kuppevelt et al., 2020).

### 2.2. Convolutional Neural Networks

Convolutional neural networks (CNN) are an influential class of deep neural networks. These networks have been mainly applied for pattern recognition in image data (e.g., Brodrick et al., 2019; Christin et al., 2019; Krizhevsky et al., 2017; Wäldchen and Mäder, 2018), but effective examples of their application for time series classification have been recently published (e.g. Zhao et al., 2017). A key component of CNNs are the so-called convolutional layers (Fig. 1b). These layers extract local features from the raw time series by applying ‘filters’. Each filter determines if a given pattern (e.g. ‘a spike’) occurs in the data and in what regions. These layers are often followed by rectified linear unit (ReLU) (or a similarly shaped function) and ‘pooling’ layers. The ReLU layers transform the summed weighted input from nodes in the convolutional layer into outputs that range from 0 to + ∞, while pooling layers reduce the dimensionality of outputs from the ReLU layer. CNNs often layer multiple instances of convolution, ReLU and pooling layers in a sequence, to build a hierarchy of increasingly abstract features. This sequence of layers is usually followed by a fully connected (or ‘dense’) layer, where each node is connected to all nodes in adjacent layers, and where classification outputs are calculated (Fig. 1b).

### 2.3. Recurrent Neural Networks

Recurrent neural networks (RNNs) are specifically designed for sequence-type input data, such as time series (Fawaz et al., 2019; Graves et al., 2013). These models are defined by inclusion of feedback loops, where the output of a layer is added to the next input and fed back into the same layer (Fig. 1c). This allows RNNs to characterize sequential patterns in the input data, but their ability to capture long term dependencies is limited due to the RNN’s tendency to prioritize signals in the short term while failing to learn long term signals (i.e., the ‘vanishing gradient problem’; Bengio et al., 1994). To overcome this problem several adaptations to the simple RNN architecture have been proposed, the most popular of which being the use of gating units (Fig. 1c, grey squares), such as ‘Long Short Term Memory’ (LSTM) and ‘Gated Recurrent Units’ (GRU) (Chung et al., 2014). Gating is a technique that helps the networks decide to either forget the current input or to remember it for future time steps, hence effectively improving the modelling of long-term dependencies (Chung et al., 2014).

### 2.4. Residual Networks

Residual networks (ResNet) are recently proposed in the context of image recognition (He et al., 2016). Basically, these networks introduce a new type of component, the ‘Residual Block’, to CNN-type models (Fig. 1d). The aim of these blocks is to allow the training of deeper models (i.e., having more hidden layers). In theory, deeper models should improve classification performances, as they allow higher levels of data abstraction. However, in practice the performances may not improve, among other things, due to the vanishing gradient problem (see above). The use of residual blocks aims to address this by forwarding the output of layers directly into layers that are several levels deeper (e.g. 2–3 layers ahead; Fig. 1d). Recently, this architecture has been applied for time series classification (Wang et al., 2017), often performing very well (Fawaz et al., 2019).

### 2.5. Inception Time Networks

Inception time networks are a very recent type of architecture, proposed specifically for time series classification (Fawaz et al., 2019). This network is an ensemble of CNN models having ResNet-type components and modules called ‘inceptions’. Inception modules ‘rework’ how convolution layers act in the networks, so that instead of being stacked sequentially, they are ordered to work on the same level in parallel (Fig. 1e). This approach allows the application of multiple filters with highly varying temporal lengths working on the same input time series. In comparison to sequential convolutional layers (as in ‘simple’ CNN) this lowers processing costs and reduces the risk of fitting noise in the data (i.e., overfitting) (Fawaz et al., 2019).

### 2.6. A standardized, accessible, deep learning modelling framework

Deep learning models can be implemented using several programming languages and specialized libraries (see Christin et al., 2019 for a review). Here, we use mcfly, a Python package for time series classification using deep learning (van Kuppevelt et al., 2020).

Mcfly utilizes TensorFlow (www.tensorflow.org) an extensively adopted machine learning library, it can make use of (but does not require) dedicated hardware (such as Graphical Processing Units: ‘GPUs’) and includes procedures for inspecting and visualizing the parameters of trained models. This software also works with both univariate and multivariate time series classification. The former refers to models using only one predictor variable (for example, when classifying species identities from bird call recordings), while the latter concerns models using two or more of these variables (for example, when time series of temperature, precipitation and wind are used to classify between the presence or absence of a phenological event). In its current version (v.3.0) mcfly allows implementing CNN, Deep convolutional LSTM (‘DeepConvLSTM’; an architecture composed of convolutional and LSTM recurrent layers), ResNet and InceptionTime architectures. Specific details about the components and structure of each architecture are given in van Kuppevelt et al. (2020).

An important feature of mcfly is that it automatizes the implementation and identification of suitable deep learning architectures and hyperparameters (i.e., ‘AutoML’; He et al., 2021). This represents an important advantage for non-experts in deep learning, as it does not require the manual assembly of the models and definition of their hyperparameters. The AutoML procedure starts by generating a set of candidate models with architectures and hyperparameters (e.g. number of layers; learning rate) selected at random from a prespecified range of values (see Fig. 2). Each candidate model is trained using a small subset of the data (data partition A*t*; Fig. 2) during a small number of epochs. After training, the performance of the candidate models is compared using a left-out validation data set (A*v*; Fig. 2). The selected candidate model (usually the best performing among candidates) is then trained on the full training data (B*t*; Fig. 2). In this step it is required to identify an optimal number of training epochs, to avoid under- or overfitting of the model. A model trained too few epochs will not capture all relevant patterns in the data, reducing predictive performance. A model trained for an excessive number of epochs might overfit, reducing its generality and ability to classify new data. There is no definitive way to identify an optimal number of training epochs, but one practical approach is through monitoring the model’s validation performance (i.e., using holdout data partition B*v*; Fig. 2). The ‘optimal’ number of training epochs is the one that provides the best validation performance. Finally, the performance of the model having an ‘optimal’ number of training epochs is evaluated using a ‘final’ test data set (T; Fig. 2), providing the best estimate of the predictive performance of the model.

**Figure 2.**
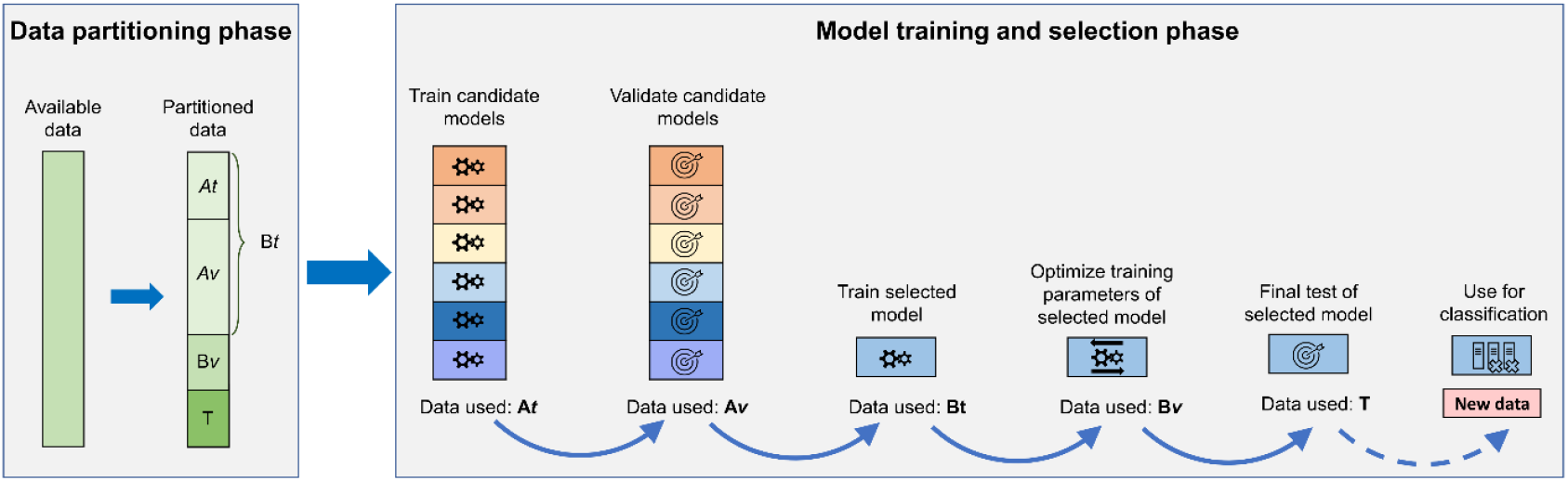
Schematic of data partitions and modelling workflow used for time series classification.

For the three case studies below, we used the same model generation and selection strategy. We had mcfly generate 20 candidate models, five for each architecture type. These models were trained during 4 epochs (using A*t*). The candidate model achieving highest performance in predicting the classes of the validation data (A*v*) was then trained on the full training data set (B*t*). For each epoch we measured training performance, as provided by mcfly (which uses the accuracy metric i.e., ‘the proportion of cases correctly classified’). The classification performance on the validation data (B*v*) was measured using the area under the receiver operating characteristic curve (AUC), a metric that is not affected by differences in the prevalence of classes and is widely used in ecology (e.g. Dyderski et al., 2018).

To identify an ‘optimal’ number of training epochs, we examined the progression of validation performance (B*v*). Models can be trained for an infinite number of epochs, so here we stopped training if no increase in validation performance was observed after 25 epochs (other thresholds could be considered, according to time resources available). Finally, the model trained with the number of epochs showing highest AUC in predicting B*v* was used to classify the test data (data set T), with performance measured using AUC.

We recorded processing time of all models from the onset of training of candidate models to the last training epoch evaluated for the selected model. This was done on two distinct systems: a ‘desktop PC’ with an Intel i7 4-Core (3.40GHz) processor and 8GB RAM and a ‘high-end workstation’ with an AMD Ryzen 9 12-Core (3.80 GHz) processor, 64 GB RAM and a NVidia RTX 2060 GPU. Because CPU- and GPU-based TensorFlow generate distinct random hyperparameters, modelling results will differ between the two computer systems. We report results and processing times for the desktop PC system. For the workstation we report processing time only. We emphasize that the timings recorded in the two systems are not directly comparable as they correspond to distinct modelling routes.

It is important to bear in mind that the modelling strategy described above aims at general applicability and further tailoring for specific classification tasks could be beneficial. For instance, with *a priori* knowledge that a specific architecture, say CNN, is best suited for the classification task at hand (see discussion section), the selection could be adjusted to generate only CNN-type candidate models. Further information about fine-tuning of mcfly model generation and selection can be found in van Kuppevelt et al., (2020).

### 2.7. The case studies

Three case studies below demonstrate the wide applicability of deep learning models for supervised time series classification, across sub-fields of ecology. Unlike most previous studies, our cases studies do not use sound or image data. Instead, they represent classification tasks that are predominantly approached by ecologists through feature-based approaches i.e., the conversion of the time series data into a set of temporally aggregated predictors of the target classes.

#### 2.7.1. Case study 1: Species identification

In this case study we predict the identity of three insect species: the olive fruit fly (*Bactrocera oleae*), the western honey bee (*Apis mellifera*), and the black fig fly (*Lonchaea aristella*) using wingbeat spectrograms (frequency series of amplitude values; Potamitis et al., 2015). *B. oleae* is an olive fruit fly pest, which if left unmanaged can lead to large economic costs worldwide (Potamitis et al., 2015). The wingbeat spectrum characteristics of these three species allow us to exemplify an ‘easy’ classification case and a ‘difficult’ classification case: while in *A. mellifera* harmonics partially overlap with those of *B. oleae*, these species show important differences in frequencies and thus constitute the ‘easy’ classification case; in contrast, *L. aristella* has a wingbeat spectrum that completely overlaps with that of *B. oleae*, representing the ‘difficult’ classification case.

We thus have three classes, each corresponding to a species ‘positive’ identity. The data does not suffer from a strong imbalance (i.e. a strong dissimilarity in the number of samples per class) and consist of 230 samples for *B. oleae*, 205 for *A. mellifera*, and 252 for *L. aristella*.

Species were identified (classified) according to their wingbeat spectrograms, which consist of frequency series of amplitudes (the predictor variable) obtained from Potamitis et al. (2015). A sample was composed of a total of 256 steps (frequencies), each step corresponding to an amplitude value for a frequency. This case study illustrates the use of these models using only one predictor variable (i.e., a single time series).

The records of species identity data and predictor variable (amplitude per frequency) were split into: data for training candidate models (∼50%; A*t*), data for validating candidate models (∼20%; A*v*), data for training the selected model (∼70%; B*t*; resulting from merging the two previous data sets), validation data for determining the number of epochs for training the selected model (∼15%; B*v*) and test data for final assessment of classification performance (∼15%; T in Fig. 2).

#### 2.7.2. Case study 2: Species distribution model

In this case study we predict the potential distribution of *Galemys pyrenaicus* (Iberian desman) using time series of environmental data. *Galemys pyrenaicus* is a vulnerable semi-aquatic species, endemic to the Iberian Peninsula and the Pyrenean Mountains. We collected distribution records from the Portuguese and Spanish atlases of mammals (Bencatel et al., 2017; Palomo et al., 2007). The data consists of 6141 UTM grid cells of 10×10 km, of which 659 record the species presence (class ‘Presence’) and 5482 its absence (class ‘Absence’).

The environmental conditions in each cell were characterized using four variables: 1) maximum temperature; 2) minimum temperature, 3) accumulated precipitation, and 4) altitude. The first three variables consist of time series of monthly values collected from CHELSA (Karger et al., 2017) spanning 1989 to 2013, totaling 300 time steps. The fourth variable was from (Fick and Hijmans, 2017) and corresponds to temporally invariant values of altitude, coded as a time series.

Species distribution data and predictors were split similarly as above with different proportions: a) A*t* ∼ 35%, b) A*v* ∼ 35%, c) B*t* ∼70%; resulting from merging A*t* and A*v*, d) B*v* ∼ 15%, and e) test data set T ∼15%. The low percentage of data used for training the candidate models in comparison to case study 1 aims to reduce computer processing time, given larger data volume.

The training and internal validation of deep learning models are sensitive to strong class imbalance (i.e., when one or several classes have a much higher number of samples). Strong class imbalance can bias models towards the prediction of majority classes (Menardi and Torelli, 2014) and reduces the reliability of performance metrics such as accuracy *sensu stricto* (i.e., the proportion of correct predictions to the total number of samples), which is used for the automated selection of candidate models in mcfly (van Kuppevelt et al., 2020). Accordingly, we balanced our data by randomly duplicating presence records and deleting absence records until a balance of ∼50:50 is obtained, which was executed using the ROSE package (Lunardon et al., 2014) for R (R Core Team, 2020). This was done for the data sets that mcfly uses for internal assessment of accuracy *s*.*s*. (A*t, Av* and B*t*, Fig. 2). Data partitioning was performed prior to balancing, to avoid inclusion of replicated cases of the same data across multiple partitions. The remaining data sets (i.e., B*v* and T) were not balanced.

#### 2.7.3. Case study 3: Phenological prediction

In this case study we predict the timing of fruiting of the *Macrolepiota procera* (Parasol mushroom) across Europe. This species produces fruiting bodies valued for human consumption (Capinha, 2019) and predicting their emergence could be useful for managing human pressure on the species and its habitats. Data is from Capinha (2019), a study employing a feature-based approach to achieve an equivalent aim. The data have two classes. One class (‘fruiting’) corresponds to locations and dates of observation of fruiting bodies of the species (from 2009 to 2015). The second class corresponds to ‘temporal pseudo-absences’, which are records in the same locations of the observation records, but with dates selected at random along the temporal range of the study (Capinha, 2019). The aim of the classification is to distinguish the meteorological conditions associated with the observation of fruiting bodies of the species from the range of meteorological conditions that are available to it.

We characterized each record using four time series: 1) mean daily temperature for the preceding 365 days, 2) daily total precipitation for the preceding 365 days, 3) latitude and 4) longitude. Time series of temperature and precipitation were extracted from the daily AGRI4CAST maps (http://agri4cast.jrc.ec.europa.eu/), at a cell resolution of 25×25 km. Geographical coordinates were coded as temporally invariant time series.

Records from 2009 to 2014 were randomly partitioned into: A*t*: 15%, A*v*: 70%, B*v*: 15%, and B*t*: 85% (merging A*t* and A*v*). Data for the year 2015 was used to evaluate the predictive performance of the final model (T), allowing comparison with the performance results achieved in Capinha (2019).

To increase the representation of the meteorological conditions occurring in the location of each observation record, the data consists of 15 pseudo-absence records per each observation record (Capinha, 2019). Similarly to the previous case study, we corrected for class imbalance by balancing the number of samples in each class using a random deletion and duplication approach (Lunardon et al., 2014). This balancing was performed for data sets A*t*, A*v* and B*t*.

## 3. Results

### 3.1. Species identification

The candidate model with greatest ability to distinguish between the spectrograms of the three insect wingbeats had an InceptionTime architecture (accuracy = 0.85; model number 15; Table 1; Fig. 3b). On the training data set this model showed a progressively increasing training accuracy with number of epochs (Fig. 3c). However, its evaluation against left-out data (B*v* data set; Fig. 2) showed that best performances were found mainly between training epoch ∼30 and ∼50 (‘validation AUC’; Fig. 3c), followed by little change. The highest validation performance was obtained after 47 training epochs. On the test data (T; Fig. 2), this model achieved an average AUC of 0.96 (Table 1), resulting from an AUC of 1 in classifying between *B. oleae* and *A. mellifera*, an AUC of 0.88 in classifying between *B. oleae* and *L. aristella* and an AUC of 1 in classifying between *A. mellifera* and *L. aristella*. Computer processing time, from the onset of candidate model training to the 72^nd^ training epoch of the selected model, took 26 minutes on a desktop PC. On the high-end workstation, a distinct modelling event took 3 minutes.

**Table 1.**
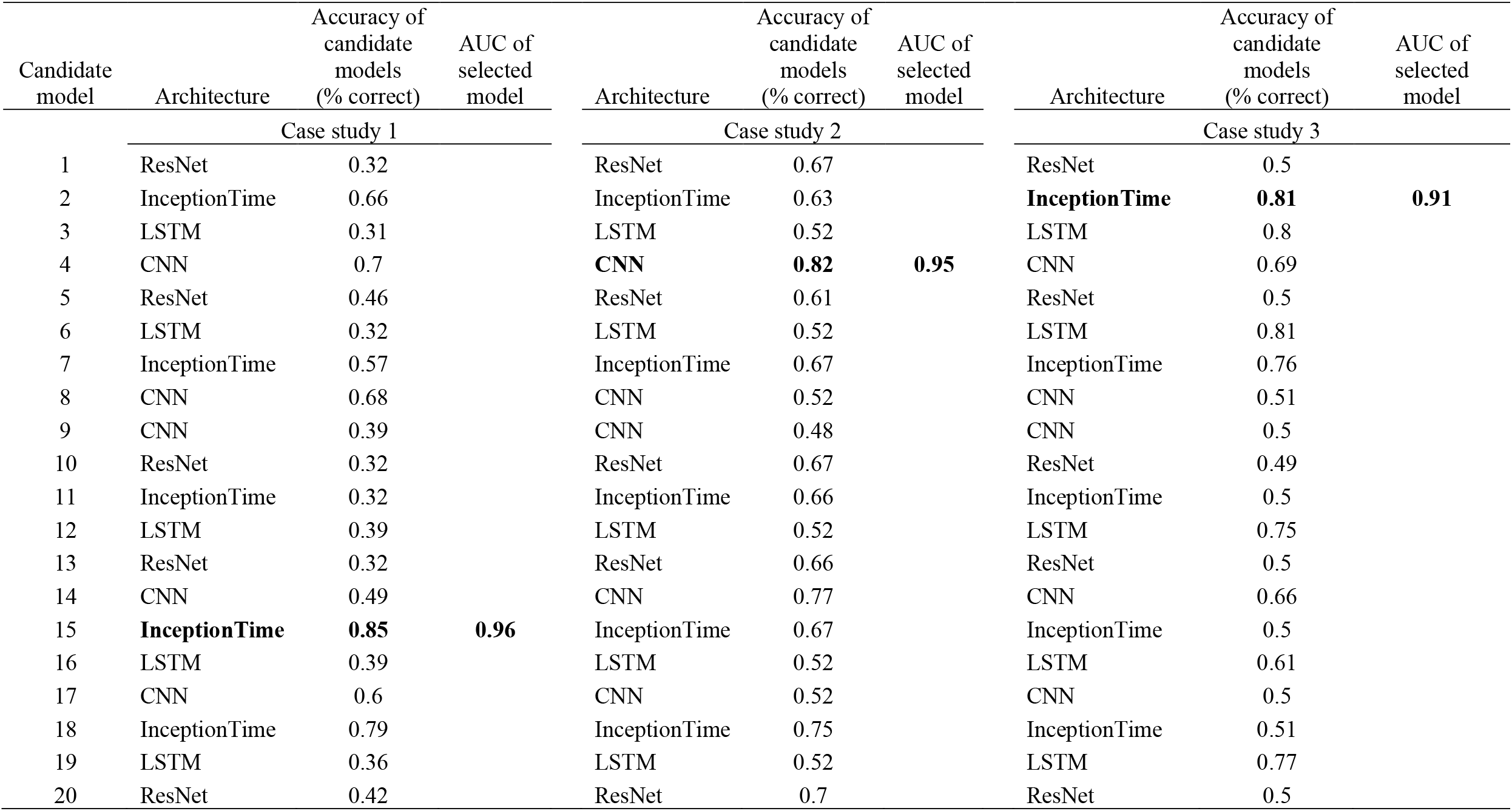
Type of architecture and accuracy of candidate models and predictive performance of selected models (bold). The accuracy of candidate models was measured using the proportion of correctly classified cases. The accuracy of selected models was measured using the area under the receiver operating characteristic curve (AUC).

**Figure 3.**
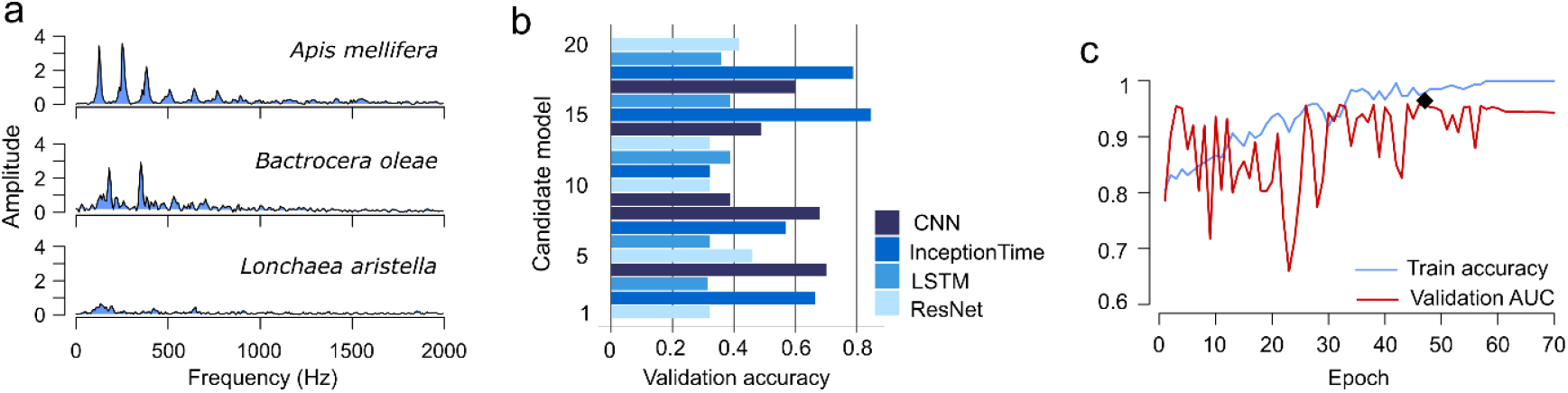
Data and results of deep learning models classifying insect species from wingbeat spectrograms. (a) Example wingbeat spectrograms for each species. (b) Validation accuracy for candidate deep learning models. (c) Training and validation curves of the selected model along time (highest validation performance is marked with a diamond symbol).

### 3.2. Species distribution model

The best performing candidate model for this case study had a CNN-type architecture (model number 4; Table 1; Fig. 4b), reaching 0.82 of validation accuracy. Using the full training data set, this model showed a decreasing trend in validation values after the ∼60^th^ epoch (B*v*; ‘validation AUC’; Fig. 4c), with highest performing classification at the 56^th^ training epoch. The model trained with this number of epochs achieved an AUC of 0.95 (Table 1) on the final test data (T). Most of northern Iberian Peninsula was predicted as suitable to *Galemys pyrenaicus*, particularly the high mountainous areas (Fig. 4e). Computer processing time took 2 hours and 49 minutes on a desktop PC. A distinct modelling event on the high-end workstation took 19 minutes.

**Figure 4.**
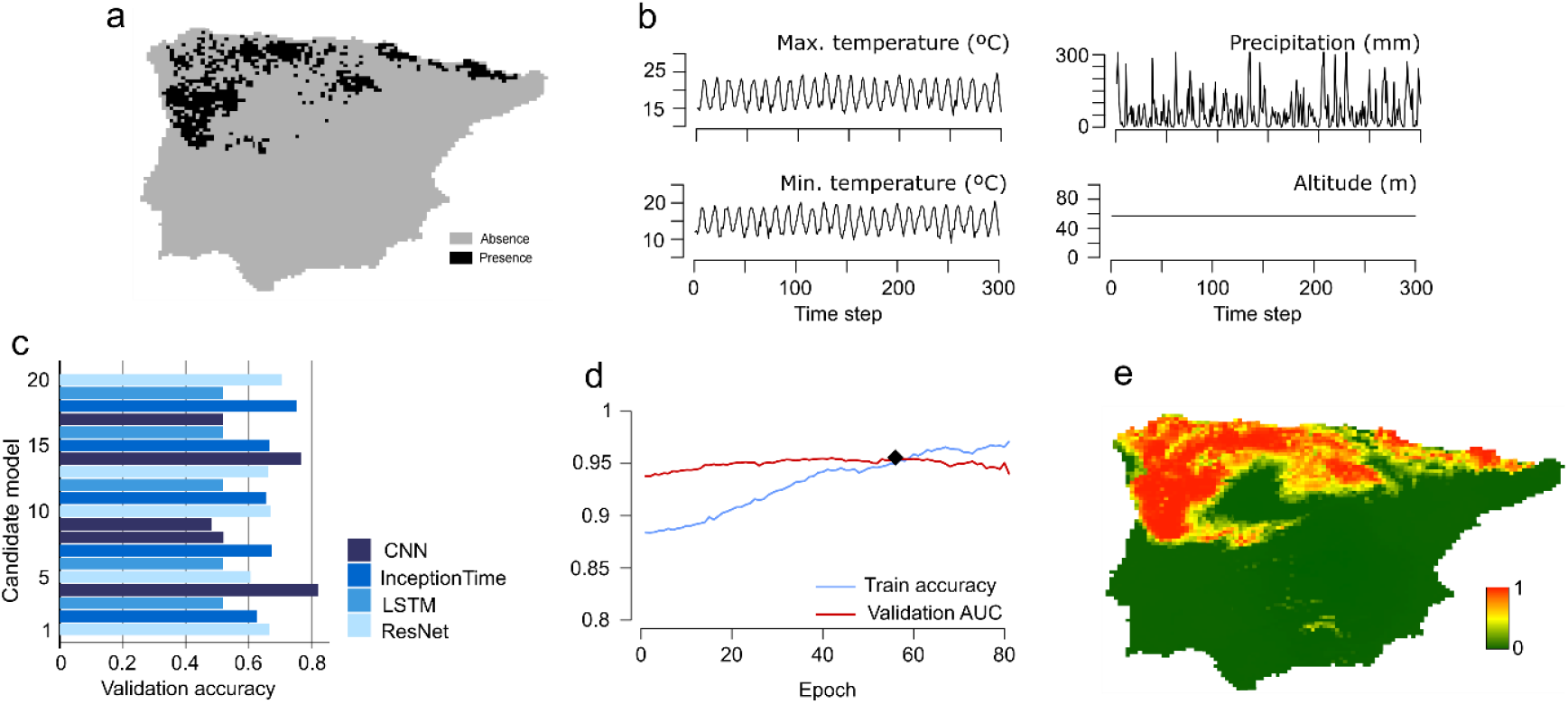
Data and results of deep learning models classifying environmental suitability for the Iberian desman. (a) Presence and absence data of the species. (b) Example of time series used as predictors. (c) Validation accuracy for candidate deep learning models. (d) Training and validation curves of the selected model along time. The diamond symbol marks the highest validation performance. (e) Environmental suitability predicted by the selected model.

### 3.3. Phenological prediction

For this case study, the selected candidate model had an InceptionTime-type of architecture (model number 2; Table 1; Fig. 5a), achieving 0.81 validation accuracy. The classification performance of this model (measured with external data; B*v*) increased only up to the 5^th^ epoch (Fig. 5b). The model trained for 5 epochs achieved an AUC of 0.91 on the final test data. The predicted probabilities of fruiting for an example site (Fig. 5c) show the ability of the model to capturing seasonal variation, with higher probabilities generally being predicted for the Autumn season, but with important inter-annual differences. Computer processing time took 10 hours and 23 minutes on a desktop PC. On a high-end workstation a distinct modelling event took 18 minutes.

**Figure 5.**
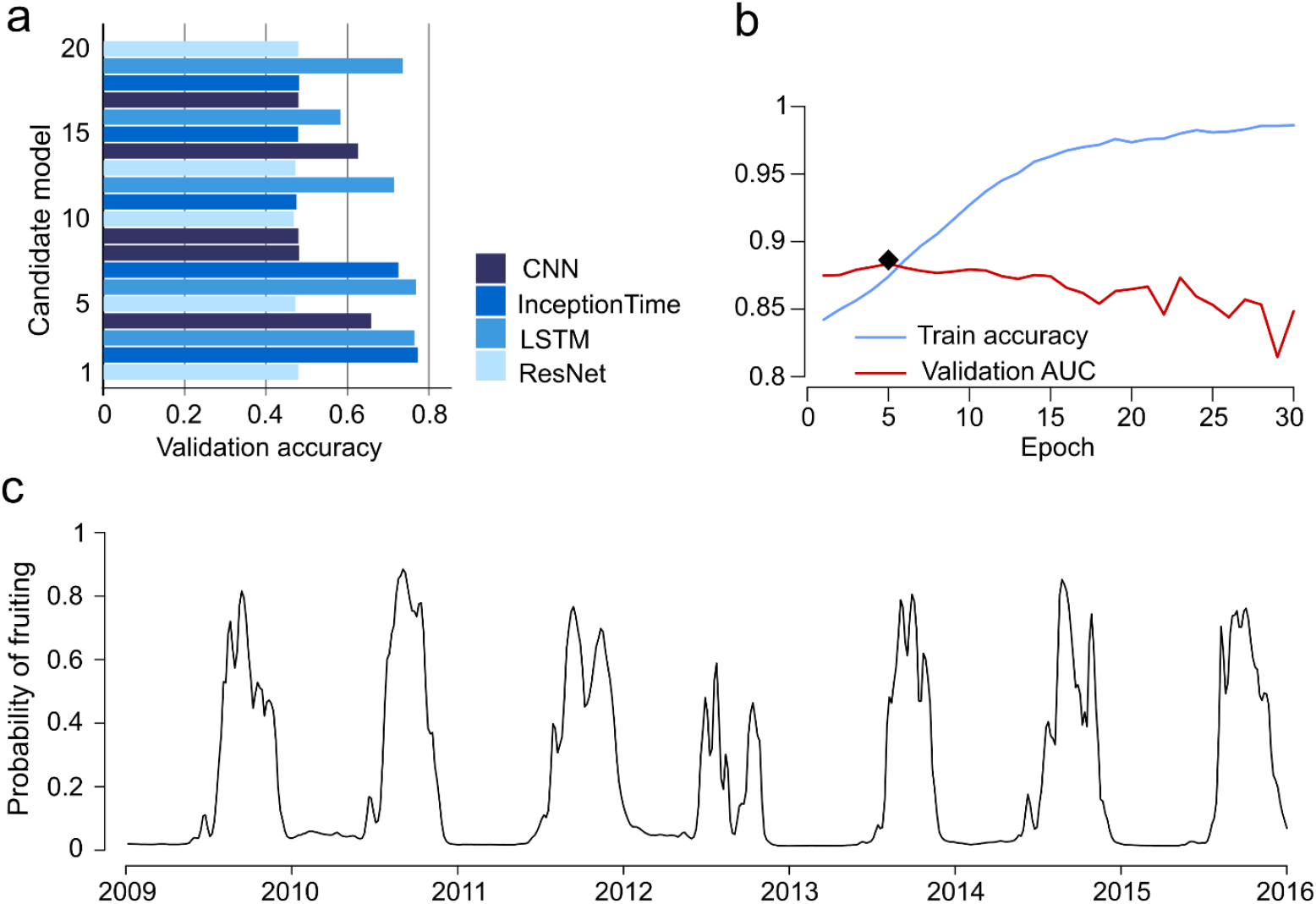
Data and results of deep learning models classifying the fruiting phenology of the parasol mushroom based on meteorological variation. (a) Validation accuracy for candidate deep learning models. (b) Training and validation curves of the selected model along time (the diamond symbol marks the highest validation performance). (c) Patterns of fruiting seasonality predicted by the selected model for an example location.

## 4. Discussion

Deep artificial neural networks are a flexible modelling technique with notable success in a range of scientific fields (LeCun et al., 2015). In ecology, the adoption of these models is still in its infancy and has been mainly directed towards image and sound recognition (Brodrick et al., 2019; Christin et al., 2019). We here introduce the use of deep learning models as a generic approach for the classification of temporal data and demonstrate how these models can be implemented and evaluated for distinct tasks across subfields of ecology.

Our case studies demonstrate the versatility and potential of deep learning for time series classification. In the first case study, an InceptionTime model performed well in distinguishing insect species based on spectrograms of their wingbeats. Potamitis et al. (2015) classified the same data using the absolute distance of spectra. They found that this distance metric achieved a good accuracy in distinguishing between *A. mellifera* and *B. oleae* (a ‘easy’ classification case) but was unable to distinguish between the latter species and *L. aristella* (a ‘difficult’ classification case). Given the use of different data partition strategies and performance metrics, the performances measured for our InceptionTime model are not fully comparable to those obtained by Potamitis et al. (2015). However, the deep learning approach correctly distinguished between all test instances of *A. mellifera* and *B. oleae* (AUC = 1) and was able to provide good classification performance for the more difficult classification case (AUC of 0.88), suggesting its superior classification ability.

In our second case study, a CNN model was used to predict the potential distribution of *Galemys pyrenaicus* in the Iberian Peninsula, based on altitude values and time series of temperature and precipitation. This model achieved a high predictive performance (AUC = 0.95) and the spatial patterns predicted are congruent with the known distribution of the species and previous predictions of Barbosa et al. (2009) - who used a Generalized Linear Model and a rich set of spatial predictors representing aspects of weather, climate, productivity, topography and geography. In addition, although the data and validation strategies used in the two studies differ, the predictive performance of the CNN, matches, or surpasses the top performing Iberian models of Barbosa et al. (2009). These results suggest the competence of the deep learning approach for species distribution modelling.

Finally, an InceptionTime model was used to predict the fruiting seasonality of *Macrolepiota procera* across Europe. The patterns of seasonality projected by the model were ecologically plausible and its predictive accuracy was high (i.e., an AUC of 0.91). This accuracy matches the one achieved by Capinha (2019), who used the same raw data. However, unlike the raw time series used by deep learning model, the study of Capinha (2019) used a feature-based classifier (boosted regression trees) and had to further process the time series data into a large set (*n*=40) of hand-crafted features reliant on domain-expertise (e.g., averaged temperatures and accumulated precipitation in previous weeks or months, growing degree days etc.). These results suggest that deep-learning models may overcome the need of processing the time series data into summary variables and of domain experts to guide this process, without sacrificing predictive accuracy.

Despite the promising results described above, the advantages of deep learning models for time series classification in ecology can only be fully appreciated with wider testing. The benchmarking of classification performances against traditional modelling approaches and the identification of factors associated with performance differences (e.g. degree of *a priori* ecological knowledge; complexity of the phenomena; volume of training data, etc.) will be of paramount importance. Deep learning models often perform in a superior manner in benchmark tests in other domains (Fawaz et al., 2019) but is also not uncommon to find other approaches providing equivalent predictions (some examples are shown in Wang et al., 2017). Given the complexity and computational demand of deep learning models (see below), it will be important for ecologists to have a better sense of when this approach is justifiable over simpler, less demanding, alternatives. Research efforts should also attempt to identify the deep learning architectures and hyperparameters that are best suited for specific ecological phenomena and data types. Thus far, classification performances from distinct deep learning typologies were compared using time series data coming from multiple domains (e.g. Fawaz et al., 2019), and the relevance of these results to ecology remains uncertain.

Differently from feature-based approaches, deep learning approaches allow classifying phenomena directly from raw time series data, a characteristic that requires ecologists to think more critically about the temporal component of the phenomena being classified. This increased relevance of the temporal dimension was, perhaps, best illustrated by using continuous climate data − instead of the usual long-term climate averages − for predicting the potential distribution of a species. However, the same sort of ‘fully’ temporally explicit approach can be exploited for virtually any ecological or biological entity or state, as long as the putative drivers have a temporal dimension. Further, the direct intake of time series data by deep learning models matches the increasing number of high frequency streams of digital data coming from distinct sources (e.g. satellite sensors, meteorological stations; Reichstein et al., 2019). The direct integration of these data into the models eliminates the need for resource consuming feature extraction procedures and is thus well-suited for operational modelling frameworks.

As for any modelling approach, the use of deep learning models for time series classification has several limitations. Two are especially prominent: the interpretability of models and computational demand. The need for interpretability of deep learning models has been well emphasized in recent literature (e.g. Reichstein et al., 2019). Unfortunately, most research on this topic has focused on models working with image data (e.g. Selvaraju et al., 2017), while much less attention has been paid to the interpretation of models for time series classification, particularly those applied to multivariate data (Shickel and Rashidi, 2020). Fortunately, a few techniques for interpreting the latter are beginning to emerge (e.g., Siddiqui et al., 2019) and, given the fast pace of deep learning research, we expect that soon deep learning models for time series classification will be no harder to interpret than those applied to image data. The challenges arising from computational demand appear harder to solve. Here we showed that ‘typical’ classification tasks can take several hours to run on a standard desktop computer. Additionally, the computational expensiveness of deep learning is expected to grow in the future (Thompson et al., 2020). To face this challenge, ecologists will likely have to move in the same direction as their fellow computer scientists and embrace faster hardware (e.g. GPUs, ‘tensor processing units’ and large-resourced cloud computing services) and scalable model implementations (e.g. distributed computing).

In conclusion, we consider that the use of deep learning for classifying temporal data in ecology could bring considerable improvements over conventional approaches. Software tools now exist that allow overcoming the implementation barrier for non-experts and state-of-the-art classification results seem a reasonable expectation for several tasks. However, only with extensive testing can the value of this approach be fully recognized. Those willing to venture through this modelling route could use the data and code we provide as a starting point.

## Data accessibility

Data and code for this study are available from: https://doi.org/10.5281/zenodo.4017750

## Acknowledgments

We thank two reviewers who helped improve this work. CC and ACH were supported by Portuguese National Funds through Fundação para a Ciência e a Tecnologia [CC: CEECIND/02037/2017, UIDB/00295/2020 and UIDP/00295/2020; ACH: PTDC/SAU-PUB/30089/2017 and GHTM-UID/Multi/04413/2013].

